# Rapid Electrochemical Biosensing of *Listeria monocytogenes* Using Rationally Designed Host-Pathogen Interface Peptides

**DOI:** 10.64898/2026.08.06.743259

**Authors:** Revital Krispin, Hen Okshtein, Yawen Song, Hadar Amartely, Zvi Hayouka, Mattan Hurevich, Nam Joon Cho, Shlomo Yitzchaik, Assaf Friedler

## Abstract

Rapid, selective detection of bacterial pathogens remains a central challenge. Here we report a label-free electrochemical biosensing approach that leverages protein-protein interaction (PPI)-derived peptides as recognition elements for rapid detection of *Listeria monocytogenes* (*LM*). The sensor design is inspired by the interaction between the *LM* virulence factor Internalin A (InlA) and the human host receptor E-cadherin (E-Cad1). Peptides derived from the InlA-binding domain of E-Cad1 were engineered as molecular recognition elements, with the E-Cad1(15-24) peptide displaying micromolar affinity and selective binding towards LM. Immobilization of these peptides on gold electrodes enabled bacterial detection by electrochemical impedance spectroscopy within 10 minutes, without labels or external signal amplification. A low peptide surface density was associated with enhanced binding-site accessibility and may facilitate multivalent interactions between the bacterial surface and the immobilized peptides. The platform produced a detectable response at experimentally tested concentrations as low as 1 CFU mL ¹ and exhibited excellent selectivity under the conditions examined. This work introduces a chemically programmable, PPI-inspired biosensing paradigm that uses a reductionist approach and could potentially be extended to other pathogen targets.

## Introduction

Rapid, selective, and sensitive pathogen detection is an important goal both in clinical point-of-care diagnostics, disease control, agriculture and food safety.^1,2^ Food-borne pathogens can contaminate food products during several stages of production.^3^ Current methods for pathogen detection can take several days, leading to products being released into the market with potential contamination.^4^ In hospitals and clinics, the gold standard for bacterial detection requires time-consuming procedures that might delay proper and rapid treatment of patients. These include culturing the bacteria, which takes days, followed by PCR and immunoassays. Altogether, these procedures often fail to provide timely results for urgent cases like sepsis. This gap highlights the urgent need for sensing technologies that provide fast, accurate, and pathogen-specific detection directly from complex samples.

*Listeria monocytogenes* (*LM)* is one of the common food-borne pathogen contaminants in the food industry.^5^ *LM* is a gram-positive bacterium that causes listeriosis, an illness that can cause symptoms ranging from mild flu-like symptoms to severe and potentially life-threatening complications like meningitis. In pregnant women, the infection can cause miscarriages.^6^ The harmful nature of *LM* is partly due to its ability to change its behaviour to withstand extreme conditions like cool environments and nutrient deficits. *LM* contamination is found mainly in the meat, fish, and processed food industries. It contaminates raw materials as well as ready-made products such as cold meat cuts, smoked fish, and even precut vegetables and fruits.^7^ *LM* takes a longer time to grow compared to other food-borne pathogens such as *S. Aureus* and *E. coli* due to it being psychrotrophic, utilizing enzymes and membrane lipids tuned for activity at lower temperatures. *LM* has more DNA damage checkpoints than other bacteria, and its stress response mechanisms extend its cell cycle to double the time of *E. coli*.^8^ *LM* is distinguished by its ability to invade and replicate in host cells. *LM* uses specific PPI to invade human cells. Once *LM* enters the human body, it penetrates non-phagocytic cells using its surface proteins from the Internalin protein family.^9^ Internalin A (InlA) is one of the proteins responsible for interactions with host-cell membrane-bound proteins, by interacting with the human E-Cadherin (E-Cad) protein. E-Cad is a part of the cadherin superfamily that is found on the membranes of epithelial cells.^8^ When InlA binds E-Cad, it induces the clustering of additional E-Cad units, creating additional interactions resulting in enveloping of the host cell membrane around *LM*. This eventually causes phagocytosis of the bacterial cell into the human cell.^8,10^ The current detection methods for *LM* require long incubation periods. This is since the samples taken have a low bacterial load, and they thus demand a long period of incubation on agar plates or liquid media for detection to be possible.^11,12,13^ The gold standard of *LM* label-free detection remains the isolation of *LM* from contaminated samples, culturing it on selective media, and performing PCR, a time-consuming process.^14,15^ Another common detection method is ELISA, which, despite being highly selective, can suffer from the same problem of being too slow.^16–18^

Electrochemical biosensors are a versatile platform for pathogen detection due to their inherent sensitivity, rapid signal transduction, and compatibility with miniaturized and portable systems.^19,20^ In these devices, the biological recognition pattern, such as antigen– antibody binding, nucleic acid hybridization, and protein-protein interactions (PPI), is converted to a measurable electrical signal. This direct translation of biochemical events into electrochemical responses enables quantitative analysis of target pathogens even at trace concentrations. The principal advantage of electrochemical transduction lies in its ability to probe interfacial changes at the electrode–electrolyte interface. When a biorecognition monolayer is immobilized on the electrode surface, the interaction with the analyte alters the interfacial charge distribution, electron-transfer kinetics, and ionic permeability. These physicochemical perturbations can be monitored using various electrochemical techniques, including amperometry, voltammetry, and EIS. EIS is a sensitive, label-free analytical technique for evaluating interfacial properties at the electrode surface. By applying an alternating current signal, EIS measures the impedance and alterations in the dielectric properties of the recognition layer. Impedance is the transfer function that comprises resistance and capacitance, corresponding to the system’s response to an alternating current (AC) signal across a range of frequencies. The dielectric properties, which typically arise from the functionalization of the electrode surface, change due to specific analyte interactions. The analyte binding changes the substrate packing and induces conformational changes at the sensor interface, thereby modulating its dielectric or electrochemical properties.^14,15,21^ Monitoring the diffusion of a redox-active species through the recognition layer, EIS quantifies the variations induced by the interactions of the bacteria and the functionalized electrode.^22–26^ EIS is a widely used technique in biosensing applications, enabling the development of highly sensitive detection methods for a diverse range of analytes, from ions to bacteria. The high sensitivity and specificity of EIS-based biosensors make them invaluable tools for various applications, such as environmental monitoring,^14^ medical diagnostics,^27–29^ and food safety analysis.^30,31^

To enable selective recognition, a highly specific molecular recognition element must be designed, immobilized on the electrode, and capable of binding the target bacterium. We have previously developed a strategy for the rational design of peptide-based molecular recognition elements based on PPI of the target moiety, and demonstrated its viability for kinase recognition.^32,33^ In the present work,^34^ we extend this approach to bacterial biosensing. Peptides derived from bacterial-interaction hotspots within cell surface proteins are employed as molecular recognition elements in the biosensor. Compared with nucleic acid–based receptors, peptides offer the advantage of enhanced hydrolytic stability.^35^

Several biosensors were developed for the rapid and specific detection of pathogenic bacteria.^35^ The antimicrobial peptide Leucin A, which targets gram-positive bacteria, was used in an EIS biosensor to detect *LM* in contaminated milk samples. The biosensor was able to detect *LM* in a low bacterial load of 10^3^ CFU/ml.^36^ In another study, a tapered optical fiber was coated with a gold nanoparticle film and was biofunctionalized with anti-InlA monoclonal antibodies. By using surface plasmon resonance (SPR) with the optical fiber, the limit of detection (LOD) was 4.3×10^3^ CFU/ml after 2.5 hours of incubation and testing.^37^ While these methods shorten detection times compared to conventional techniques, further shortening is still essential. Here, we describe the development of a biosensor based on the interaction between the bacterial InlA protein and the cellular E-Cad1 protein.^8,10^ Three Peptides derived from the InlA-binding hot spot of E-Cad1 were designed as molecular recognition elements and immobilized to gold electrodes. The E-Cad1 15-24 peptide enabled the detection of *LM* at concentrations as low as 1 CFU/mL within minutes. By leveraging the bacterial invasion mechanism of *LM*, this approach combines high selectivity, operational simplicity, and scalability, offering a powerful alternative to conventional methods for food safety and clinical diagnostics.^6,35^

## Results

### Rational design of InlA-binding peptides from E-Cad1 as recognition elements for *LM* biosensors

The interaction between InlA and E-Cad1 is mediated by the leucine-rich repeat (LRR) domain of InlA, which forms a highly complementary interface with the distal domain of E-Cad ^10^ (Fig. 1^38^). The InlA – E-Cad1 interface has a “ball-in-joint” conformation,^10^ where E-Cad1 interacts mainly with the β-sheet structure of InlA. Most LRR β-strands of InlA contribute to the interaction, except for LRR3 and LRR10. In E-Cad1, the tightest InlA binding regions include β-strands a and b and the intermediate loop (ab loop) that spans residues 3-32 of E-Cad1 (residue numbering is based on the PDB 1O6S^10^ structure). Val3 and Ile4 of strand a form a hydrophobic contact with aromatic residues in LRR12 and LRR15 of InlA. The ab loop contains Pro16 and Pro18, which adopt a cis-conformation. Pro18 interacts with Phe150 of InlA. Ca^+2^ ions bridge Glu326 of InlA and Asp29 of E-Cad1, while Cl^−^ ions mediate electrostatic interactions between charged residues on both proteins.

**Figure 1.**
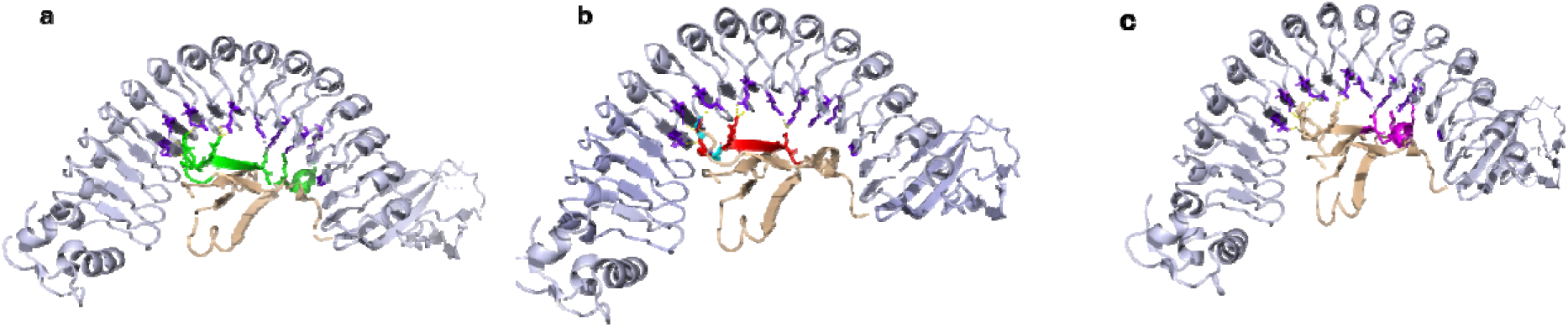
Designed peptides from E-Cad1. (Light blue – InlA, beige – E-Cad1). **a** – E-Cad1 11-31 (green), **b** – E-Cad1 15-24 (red) (Pro 16 and 18 in cyan), **c** – E-Cad1 22-31 (pink). The figure was made using PyMol^38^.

To design peptides that specifically bind *LM* and can be used as molecular recognition elements for its detection, we used the crystal structure of the InlA–E-Cad1 complex (PDB ID: 1O6S).^10^ We designed three peptides based on the E-Cad1 binding interfaces of InlA E-Cad1, focusing on regions with numerous electrostatic and hydrophobic contacts. The major interaction hotspot is around the loop containing Pro16 and Pro18, extending into the adjacent β-strand and α-helix (residues 15-31) of E-Cad1. This region includes electrostatic interactions between Lys19 of E-Cad1 and Glu255 of InlA (LRR9), and polar contacts involving Gln23 of E-Cad1 with Asn259 and Lys301 of InlA (LRR9 and LRR11). Further interactions are formed by Lys25 and Lys30 of E-Cad1 with Glu323 and Glu326 of InlA, respectively. All contacts in this region are within a distance of 3.2 Å. Based on this, the first peptide we designed was E-Cad1 11-31, which spans the loop, β-strand, and α-helix of residues 15-31, encompassing all major interacting residues (Fig. 1a). Additional designed peptides include E-cad1 15-24, which contain Pro16 and Pro18 (Fig. 1b), and E-Cad1 22-31, covering the loop and α-helix (Fig. 1c). Sequences of the designed peptides are listed in Table 1. The peptides were synthesized and labelled either 5,6-Carboxy Fluorescein (FAM) or Alpha Lipoic Acid (LPA) at their N-termini. For the analytical data, see Table S1.

**Table 1:**
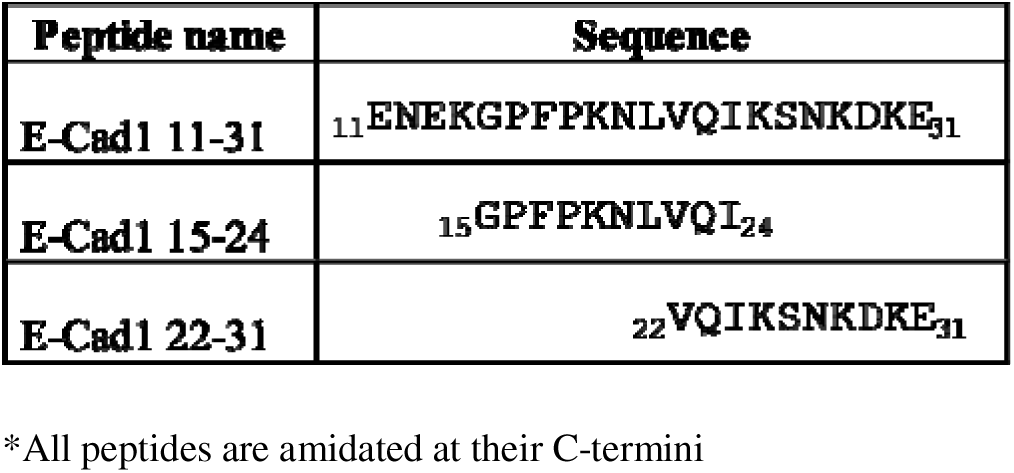
Sequences of the designed peptides*.

### E-Cad1 11-31 and E-Cad1 15-24 selectively bind *LM* with low micromolar affinity

The binding of the peptides (at concentrations of 0.1 to 25 μM) to *LM* was tested using flow cytometry. Binding to E. coli (Gram-negative) and MRSA (Gram-positive) was also studied to test the selectivity of the peptides to *LM*. At a concentration of 100 nM, E-Cad1 11-31 and E-Cad1 15-24 exhibited strong and specific binding to *LM*, with minimal binding to E. coli and MRSA. In contrast, E-Cad1 22-31 showed no detectable binding to any of the tested bacteria (Fig.2). Gating boundaries were calibrated against unexposed bacterial controls to account for their autofluorescence, revealing a multi-log shift exclusively upon specific peptide binding post-incubation. (Fig. S1)

**Figure 2:**
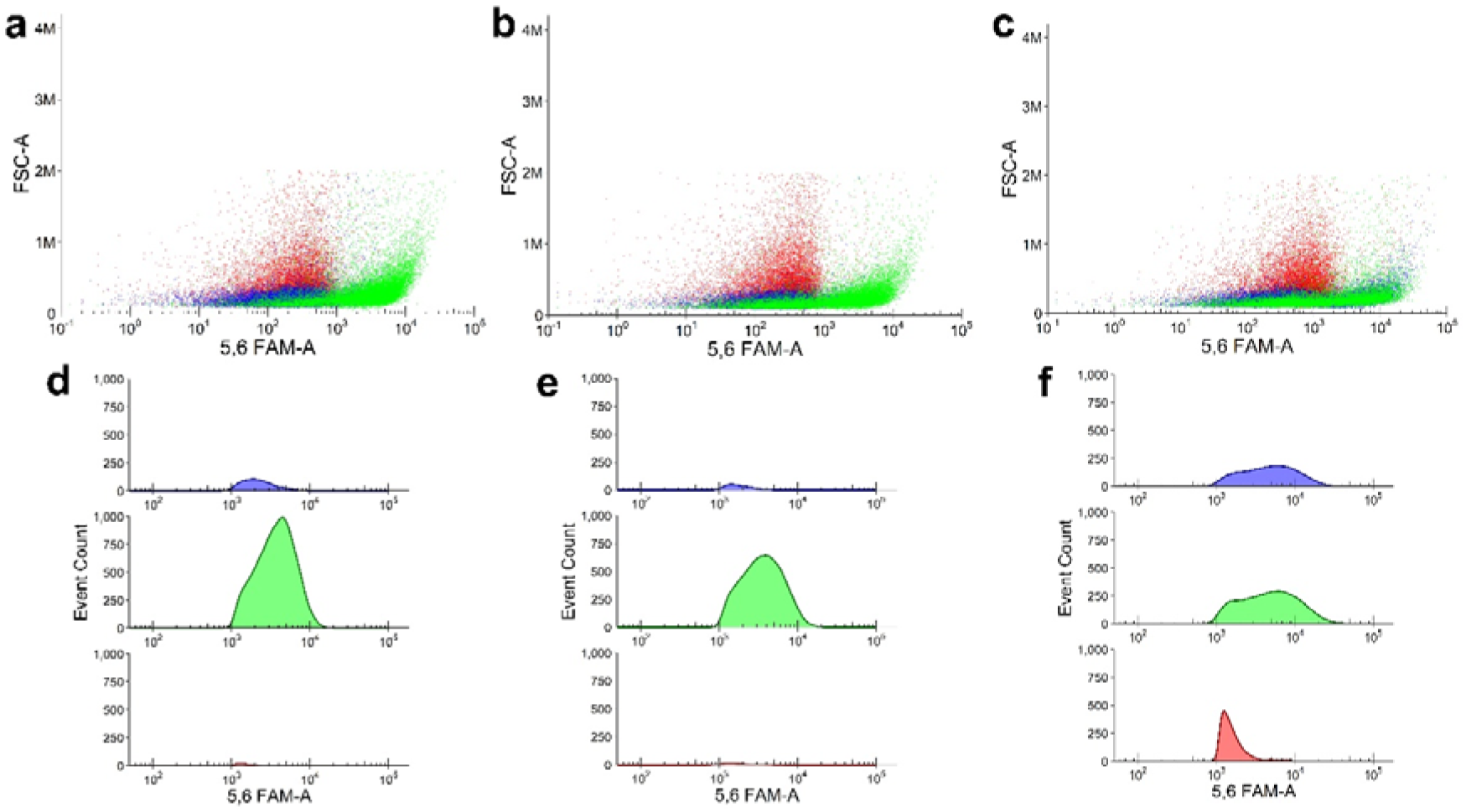
Binding of fluorescein-labeled E-Cad1 peptides to *LM* (Green) compared to *E. coli* (Blue) and *MRSA* (Red) analyzed using flow cytometry. **a-c** Dot plots of fluorescence intensity for each E-Cad1 peptide at 0.1 µM concentration, showing their specificity toward *LM* compared to *E. coli* and *MRSA.* Shown are **a** 11-31, **b** 15-24, **c** 22-31. **d-f** Event count histograms of gated fluorescence intensity for E-Cad1 peptides **d** 11-31, **e** 15-24, and **f** 22-31 at 0.1 µM concentration, displaying the number of binding events for *LM* compared to the other bacteria.

The binding curves of E-Cad1 11-31 and E-Cad1 15-24 were fit to the Hill equation, revealing cooperative binding. The apparent binding constants to *LM* were determined to be *K*_D_ = 7.4±0.3 μM for E-Cad1 11-31 and *K*_D_ = 5.7 ± 0.9 μM for E-Cad1 15-24 (Fig. 3). Based on these results, E-Cad1 11-31 and E-Cad1 15-24 were selected for electrochemical sensing experiments (Fig. 3).

**Figure 3.**
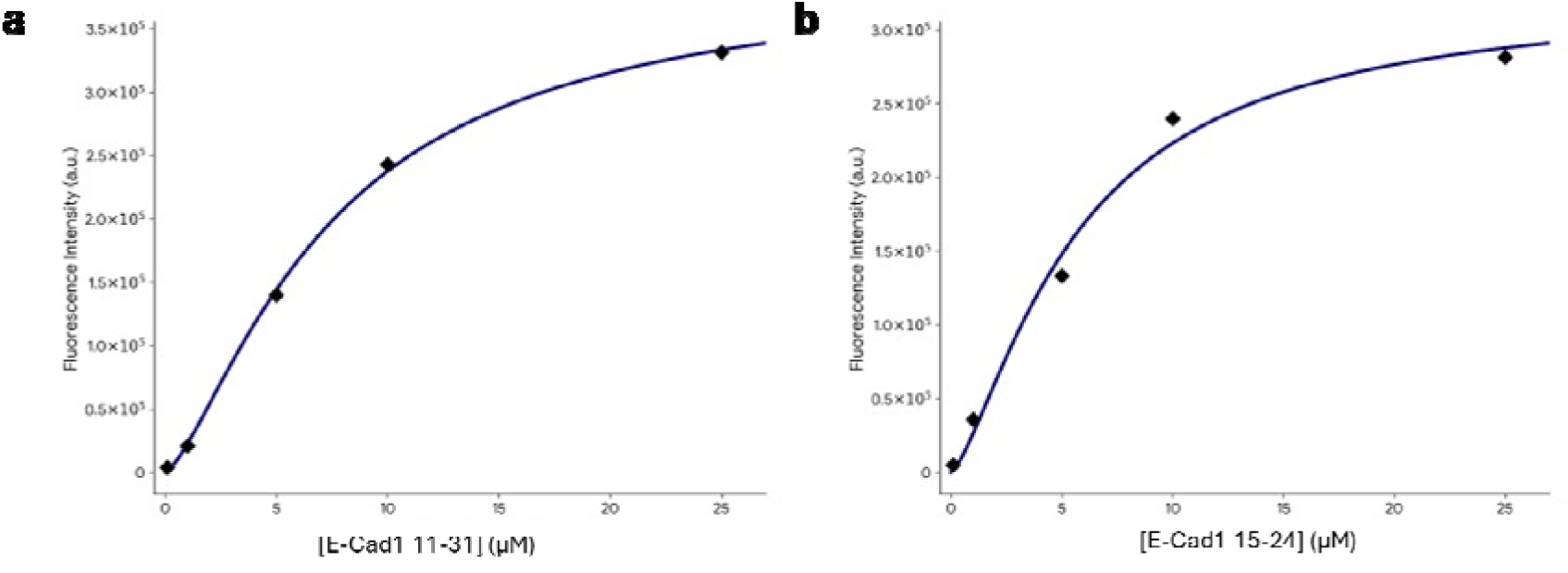
Mean fluorescence intensities at each concentration for E-Cad1 peptides **a** 11-31 and **b** 15-24, fit to the Hill equation, demonstrating the cooperative binding to *LM.* The standard deviation in each experimental data point is 0.5%-1.5% of the value.

**Figure 4.**
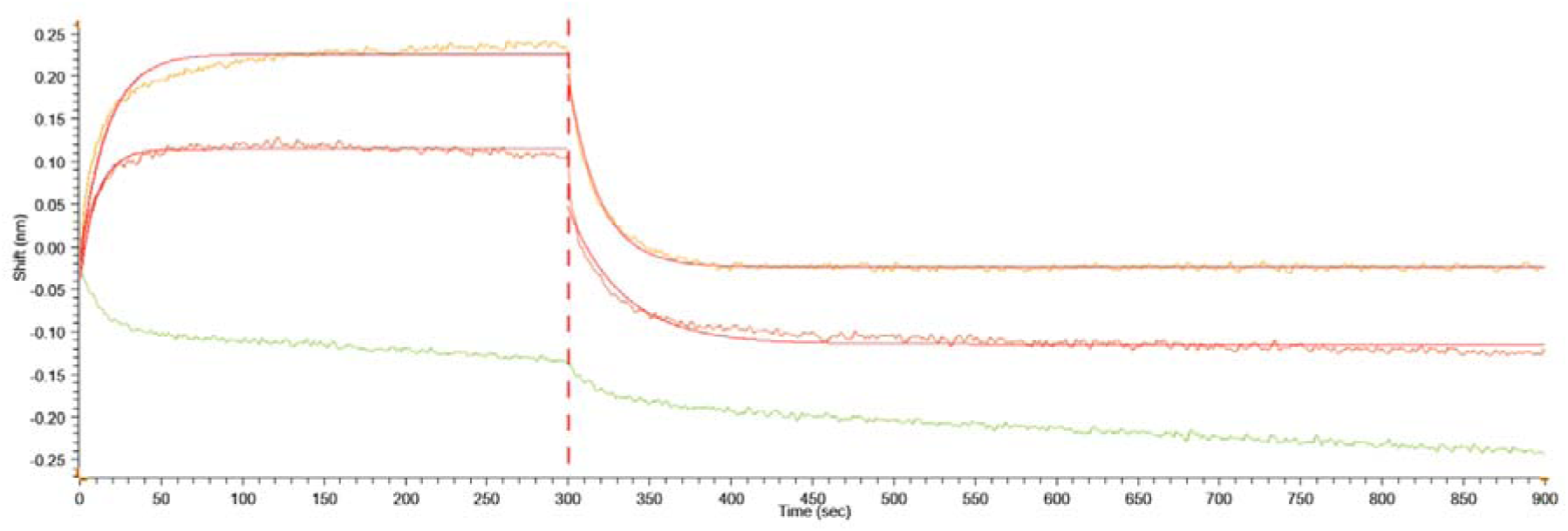
BLI Sensorgrams display the association and dissociation phases of E-Cad1 11-31 (Red), E-Cad1 15-24 (Orange), and E-Cad1 22-31 (Green) at a concentration of 10 µM upon incubation with the InlA-bound sensor.

### Biolayer Interferometry (BLI) Binding Assays

To confirm that the bacterial binding to the peptides is indeed mediated by the interaction with InlA, we performed Biolayer Interferometry (BLI) binding studies between the E-Cad1 peptides and the InlA protein. His-tagged InlA was immobilized onto the Ni biosensor surface, and binding was evaluated to a range of peptide concentrations for the E-Cad1 peptides. Both E-Cad1 11-31 and E-Cad1 15-24 bound InlA, with K_D_ of 9 ± 6 µM for E-Cad1 11-31 and 40 ± 20 µM for E-Cad1 15-24. E-Cad1 22-31 did not bind InlA. (Figure).

### Surface Characterization of Peptide-Modified Electrodes

Following peptide anchoring to **AuE**, the surface characterization of E-Cad1 15-24 peptide-modified **AuE** (**AuE**–E-Cad1 15-24) was carried out by X-ray photoelectron spectroscopy (XPS) and variable angle spectroscopic ellipsometry (VASE) (Fig. S). The XPS spectra, specifically the N1s spectra, exhibited a distinct peak at 401.1 eV, which is characteristic of the amide group, supporting the successful peptide modification by the presence of amide bonds on the modified **AuE**. The modified **AuE** surface thickness was characterized by rocking angle XPS and VASE, with an increase of approximately 20 Å (Fig. S). Reductive desorption measurements yielded a surface coverage of 3.35×10^13^ molecules/cm² (Fig. S1) with a calculated footprint of 323 Å². The peptide length, as estimated from the size of the corresponding protein segment in the crystal structure,^10^ is 22.6 Å. The addition of Lipoic acid is estimated to increase its length to approximately 31.6 Å.

The observed results are in line with the literature, as the monolayer thickness and peptide surface density results indicate that the E-Cad1 15-24 peptides on the **AuE** surface adopt a tilted orientation. Based on the ratio of measured to theoretical thickness (cos θ = 20 Å/31.6 Å), the peptide layer has a tilt angle of approximately 50° from the surface normal. The observed thickness for peptides with the same length is in the range 15–25 Å, and tilt angles of 40–60°, in line with the literature.^39,40^ The measured tilt angle and surface coverage indicate that the E-Cad1 15-24 layer forms a disordered, low-density monolayer rather than a well-packed SAM. The different packing density can be advantageous for biosensing applications: the large molecular footprint and tilted configuration maximize solvent and analyte accessibility and ensure that a substantial fraction of each peptide remains exposed and available for effective interactions with the bacterial cell surface.

Following the characterization of the peptide monolayer on **AuEs**, the surface electronic properties were characterized using contact potential difference (CPD) measurements. The **AuEs** exhibited a decrease in work function (Φ) of ΔΦ = −0.781 eV relative to the bare Au surface. Subsequent exposure to *LM* led to an additional decrease, with a total ΔΦ of −0.881 eV. These changes may correlate with the positively charged lysine residues in the peptide and the two-stage modification of the **AuEs**.

### Selective and sensitive *LM* detection using EIS

Bacterial detection was performed using EIS (Fig. 5). *LM* recognition by the biosensor was evaluated by exposing the **AuE**s modified with E-Cad1 11-31, E-Cad1 15-24, and E-Cad1 22-31 to 10^8^ CFU/ml *LM* for 10 min. E-Cad1 15-24 showed the highest increase in R_CT_, indicating stronger binding of *LM* bacterial cells to the electrode surface (Fig. 6a). E-Cad1 11-31 also showed binding, represented by a decrease in R_CT_ (Fig. 6b). This phenomenon will be discussed in a follow-up study. The E-Cad1 22-31-based electrode showed no binding of *LM* (Fig. 6c). These results demonstrate the rapid response time of the biosensor.

**Figure 5.**
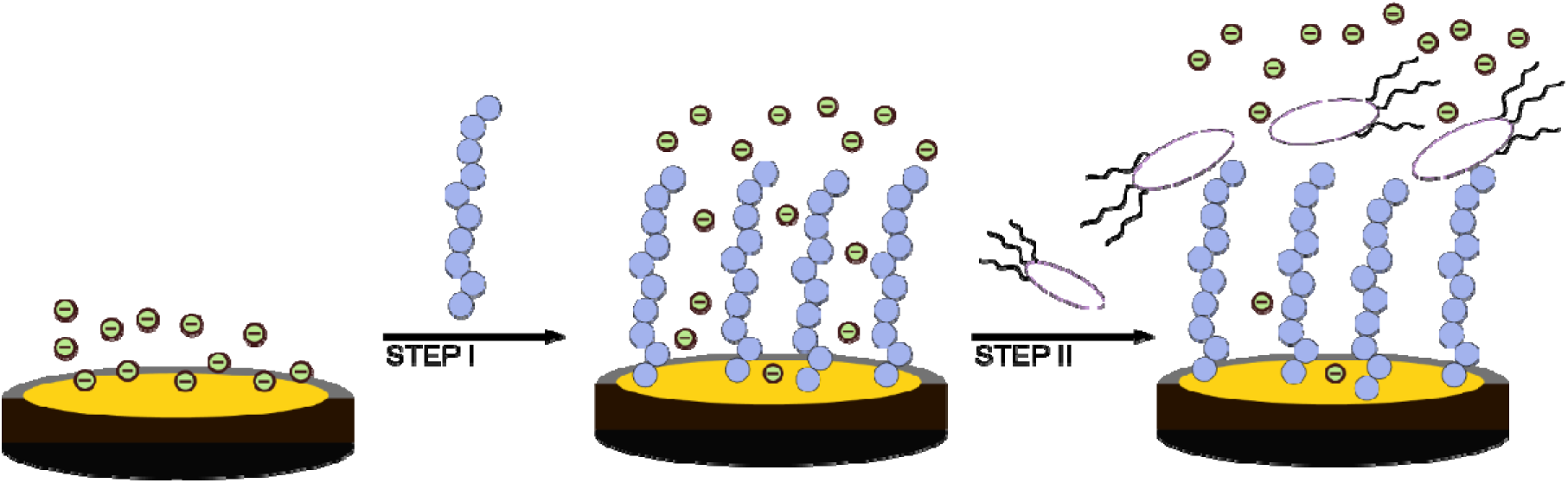
Schematic description of the biosensor assembly and detection. The sensor is based on the ability of EIS to quantify the resistance to charge transfer (R_CT_) of the peptide-modified gold electrodes following the interaction with the bacteria. Assembly steps: **step I,** peptides assembly; **step II,** exposure to bacteria.

**Figure 6.**
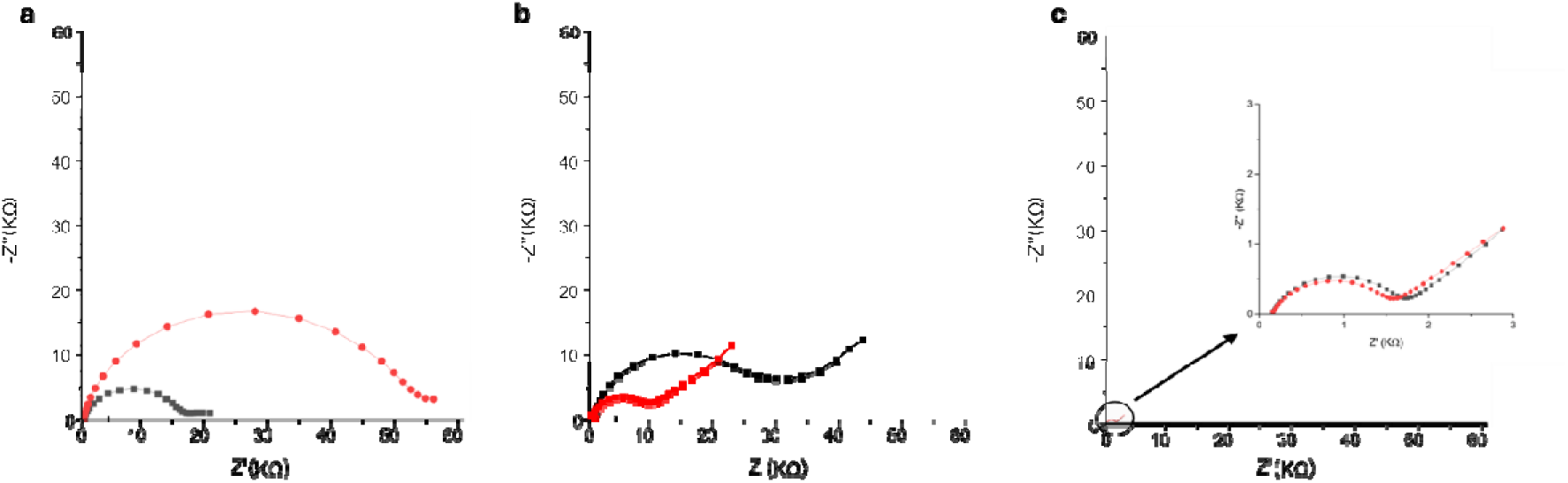
The Nyquist plot represents 3 different E-Cad1 modified **AuEs** exposed to10^8^ CFU/ml *LM*, the black plot represents the peptide-modified **AuE** before the exposure to *LM, and* the red plot represents the peptide-**AuE** after the exposure to *LM*. (**a)** The E-Cad1 15-24 shows binding to *LM* (positive response), (**b)** the E-Cad1 11-31 peptide shows binding to *LM* in a different mechanism (negative response), (**c)** the E-Cad1 22-31 shows no *LM* binding.

To confirm that the impedimetric response of E-Cad1 15-24, is driven specifically by binding to the InlA protein, the functionalized AuEs were exposed directly to recombinant InlA. The E-Cad1 15-24 modified electrode exhibited a distinct increase in R_CT_ upon incubation with InlA (Fig.7).

**Figure 7.**
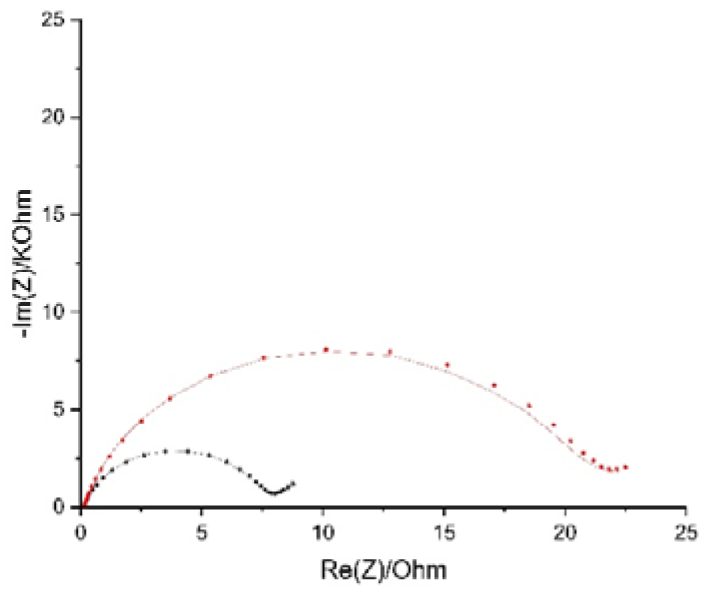
Nyquist plot representing the E-Cad1 15-24 modified **AuE** exposed to recombinant InlA protein. Baseline impedance of the peptide-modified **AuE** before exposure (black). The peptide-AuE after exposure to isolated InlA (red).

Based on the recognition results, we selected E-Cad1 15-24 as the lead peptide and further characterized the selectivity of the **AuEs** modified with this peptide to *LM* by comparing it to the response to *MRSA* and *E. coli*, at a bacterial load of 10 CFU/ml for 10 min. The biosensor demonstrated negligible binding to *MRSA* and *E. Coli* (Figure 8a), demonstrating the selectivity of the E-Cad1 15-24 peptide toward *LM* and rapid detection time. Fluorescence microscopy using Nile Red staining also confirmed the specific binding of *LM* to E-Cad1 15-24 peptide-functionalized gold surfaces, whereas control strains, *MRSA* and *E. coli*, showed negligible binding (Fig. S3).

**Figure 8.**
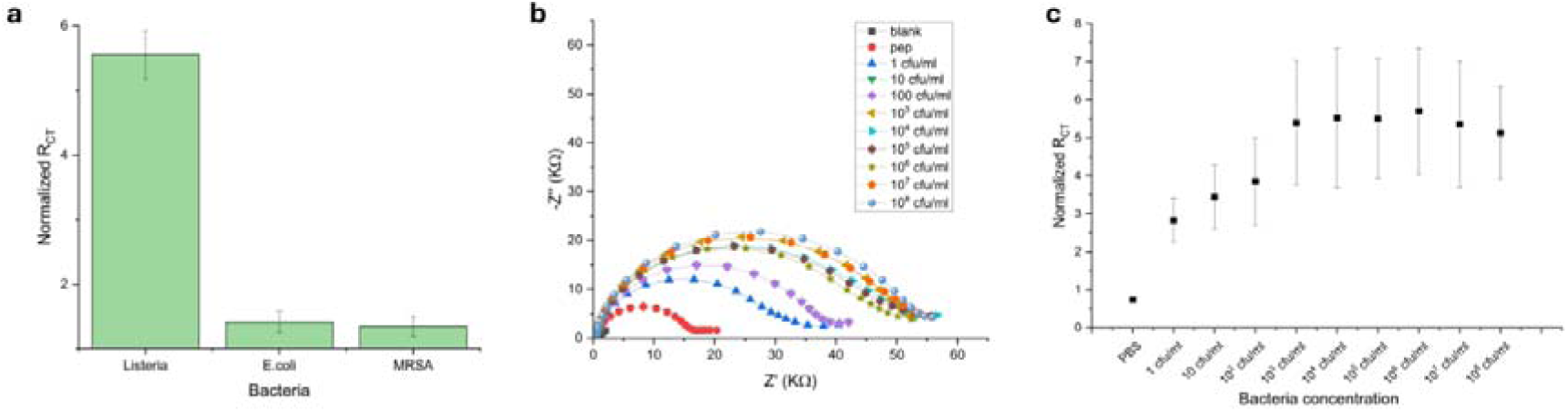
(**a**) Dose response of E-Cad1 15-24 – LPA modified **AuEs** upon exposure to increasing concentrations of *LM*. The graph shows detectable response down to as low as 1 CFU/ml and saturation at 10^3^ CFU/ml. (**b**) Nyquist plots corresponding with the dose-response graph. (**c**) Normalized R_CT_ values of **AuE**-E-Cad1 15-24 after exposure to bacteria demonstrating selectivity towards *LM* over *E. coli* and *MRSA*.

Dose-response experiments were used to evaluate the sensitivity of the biosensor. The E-Cad1 15-24 modified electrode was exposed to increasing bacterial loads of *LM*, ranging from 1 to 10^8^ CFU/ml (Fig. 8b,c), using five replicate electrodes to establish the biosensor limit of detection (LOD). AuE-E-Cad1 15-24 showed detectable response down to as low as 1 CFU/ml and saturation from 10^3^ CFU/ml. The LOD was calculated to be 0.5 CFU/ml (see materials and methods). The linear response range is the range where early detection of LM is required. The saturated region could also be observed by fluorescence microscopy. (Fig. S3)

## Discussion

In this study, we present a highly sensitive and selective peptide-based biosensor for the detection of *LM,* where the molecular recognition element is based on specific PPI of the bacteria^8,10^, enabling rapid and label-free detection. Our rational design was based on the interaction between InlA and E-Cad1. Of the three designed peptides based on this interface, E-Cad1 15-24 bound *LM* tightly and mediated specific sensing of *LM*, while the partly overlapping peptide E-Cad1 22-31 did not show any detectable binding or sensing. Within the parent E-Cad1 protein, the E-Cad1 15-24 sequence has a structure of a loop and a beta strand, which contains Pro16 and Pro18 (Fig. 9a). These residues are crucial for mediating the interaction of E-Cad1 with InlA.^8,10,41^ Substituting Pro16 with Glu, a residue naturally found in the same position in murine E-Cadherin, is sufficient to prevent InlA binding and abolish bacterial invasion^41^. In contrast, E-Cad1 22-31 contains the end of the beta strand, continues into a loop followed by an alpha helical structure, and lacks the Pro residues (Fig. 9b). This explains the molecular basis for the differential activity of the two peptides. Sensing by E-Cad1 15-24 indeed relies on the precise recognition by the InlA - binding motif in E-Cad1(Fig. 9c).

**Figure 9.**
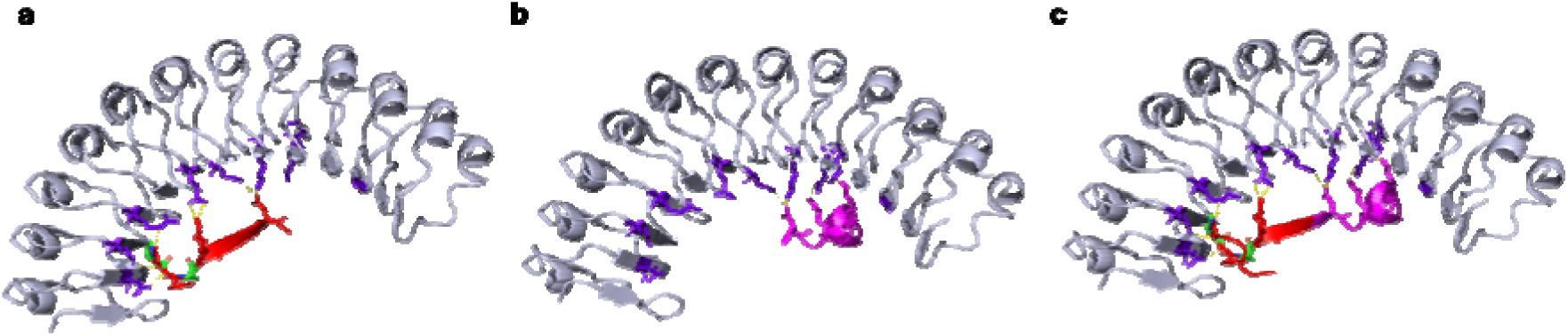
Secondary structures of the E-Cad1 peptides. InlA is in light blue with the peptide-binding residues in purple. **(a)** E-Cad1 15-24, in red, forms a beta strand, with the Pro residues in green mediating the interaction. **(b)** E-Cad1 22-31, in magenta, adopts an alpha helical structure. **(c)** E-Cad1 11-31, combining both peptides (residues 11-24 in red and residues 25-31 in pink).

The specific binding and recognition of *LM* with our biosensor is driven by the PPI between the bacterial InlA protein and the Ecad1 11-31 and 15-24 peptides. InlA is expressed only on the surface of *LM,* and the interaction of *LM* with E-Cad1 is thus unique to this bacterium.^42,43^ BLI studies with purified InlA protein confirmed that E-Cad1 11-31 and 15-24 indeed bind LM by binding directly to InlA, with micromolar affinity. The protein was bound to the sensor surface, and the peptide was in solution because of the solubility limits of the protein, which prevented achieving higher concentrations. Furthermore, the direct impedimetric response of the E-Cad1 15-24 modified electrode to recombinant InlA (Fig. 7) confirms that this specific biorecognition is successfully retained on the biosensor surface. Together, these results prove that the sensing is driven by the targeted InlA–E-Cad1 interaction.

The high sensitivity of our system is partly due to the low peptide density on the electrode surface. The low peptide packing density allows for higher conformational freedom, consistent with the observed peptide 50° tilt angle. The existence of multiple InlA proteins on the bacterial surface enables simultaneous interactions of several InlA proteins with several immobilized peptides, increasing the resistance to charge transfer and the impedimetric response through multivalent protein-peptide interactions. This behavior is consistent with previously reported electrochemical aptamer and peptide-based biosensors, where low packing densities yield higher sensitivity than densely packed self-assembled monolayers. This is because decreased molecular crowding permits larger binding-induced conformational changes and more efficient signal transduction.^44,45^

*LM* is a rod-shaped bacterium with typical dimensions of 0.5-0.8 µm in width and 2-3 µm in length, and a projected footprint of approximately 2 µm². The **AuE**s have a surface area of 2 mm^2^ per electrode (3.14 × 10 µm²) and can theoretically accommodate about 10 bacteria in a close-packed monolayer. However, under the experimental conditions, only part of the theoretical maximum, about 10³ cells (Fig. 8c), bound to the electrode according to our calculations. Based on the reductive desorption (RD), the calculated peptide surface density of about 10¹³ molecules/cm² (Fig. S1), the **AuE** monolayer has about 10¹¹ E-Cad1 (15-24) peptides. Immunogold labeling demonstrates that each *LM* displays about 10² surface accessible InlA proteins ^46^ The dose-response (DR) data present a concentration-dependent increase in normalized R_CT_ from 1 to 10³ CFU/mL, followed by signal saturation at higher bacterial loads. Under 10³ CFU/mL, given that each bacterium can engage tens of peptides per binding event through multivalent InlA–peptide interactions, the total peptide population vastly exceeds the binding demand even at high bacterial loads, indicating that saturation is governed by electrochemical blocking of the interfacial region rather than peptide depletion. At 10³ CFU/mL, the accumulated bacterial load creates sufficient InlA - peptide complexes and a negatively charged surface that induces electrostatic repulsion, inhibiting further bacterium binding, and the impedance response approaches a plateau. This indicates that decreasing the peptide density of the surface is crucial for optimizing the device response to the bacteria.

The binding affinities of the E-Cad1 peptides to InlA are in the micromolar range, as shown both at the protein level and at the bacterial level. Still, a much higher sensitivity was observed in the impedimetric sensing. This is because a single binding event is amplified by the large area that the bacteria cover on the peptide layer, resulting in a stronger signal.

Our platform offers a distinct performance advantage over existing detection methods and strategies. The E-Cad1 15-24 biosensor demonstrated high selectivity toward LM and produced a detectable response at experimentally tested concentrations as low as 1 CFU mL ¹. An LOD of 0.5 CFU mL ¹ was estimated from the calibration response using the 3σ method. This value represents a statistical extrapolation and was not directly validated by measurements performed at 0.5 CFU mL ¹. Nevertheless, the extrapolated LOD is numerically lower than those reported for several antibody- and antimicrobial peptide-based detection methods. The Leucin A peptide-based sensor has a LOD of 10^3^ CFU/ml and exhibits cross-reactivity with other Gram-positive bacteria.^36^ Fiber-optic immunosensors require 2.5 hours of incubation and have a LOD of 4.3×10^3^ CFU/ml.^37^ DNA aptamers that capture *LM* typically involve multistep processes.^47^ For example, using biotinylated DNA aptamers conjugated to magnetic beads that capture and concentrate the bacteria resulted in an LOD of 10^2^ CFU/ml. However, this approach relies on a downstream qPCR process that requires hours to analyze. Our peptide biosensor offers a robust and operationally simple alternative, taking 10 minutes of incubation on the AuE. The E-Cad1 15-24 peptide directly mimics the *LM* target protein during its invasion, and this straightforward recognition, based on its natural target, results in a much lower LOD without the need for complex antibodies or aptamers or the need for labeling.

The E-Cad1 11-31 peptide, which contains 21 residues, displayed a unique electrochemical behavior compared to its shorter and more active derivative E-Cad1 15-24. Although the E-Cad1 11-31 peptide successfully bound *LM*, it generated an inverse impedimetric response compared to the positive signal of the E-Cad1 15-24. The signal inversion indicates that the interface of E-Cad1 11-31 with *LM* is different from that of E-Cad1 15-24. Given its counterintuitive behavior, a detailed characterization will be discussed in a follow-up study.

In summary, this study establishes that rationally designed peptides derived from the host-pathogen interfaces can serve as highly effective biorecognition elements for impedimetric biosensors. The E-Cad1 15-24 biosensor provides a rapid, sensitive, selective, and robust platform for *LM* detection, achieving an LOD significantly lower than previously described antibody- and antimicrobial peptide-based detection methods. By exploiting the specific InlA – E-Cad1 PPI, the sensor mimics the native interaction of the bacterial InlA protein using short synthetic peptides, offering high robustness and rapid results. Consequently, this platform offers a promising, scalable solution for real-time pathogen monitoring in the food industry and clinical diagnostics.

## Methods

### Peptide Synthesis and Characterization

The peptides were synthesized on a Rink Amide resin utilizing the Liberty Blue Microwave-Assisted Peptide Synthesizer^48^ (CEM) using standard Fmoc chemistry, HATU/DIEA used as coupling reagents. The peptides were cleaved from the resin using 95:2.5:2.5 Tri-Fluoro Acetic Acid (TFA): Triple distilled water (TDW): Tri-Isopropyl-Silane (TIPS) solution. Before cleavage, some of the resin was labelled with either FAM or LPA at the N-terminus as described.^49^ The resin was added to an overnight labelling cocktail made of 1.7 ml N-Methyl-2-Pyrrolidone (NMP), 200 μl Dichloromethane (DCM), 46 μl DIEA, 42 μl N,N′-diisopropylcarbodiimide (DIC), 36 mg HoBt, and 100 mg of LPA or FAM. 100mg of the peptidyl-resin was added to the cocktail and was left to shake overnight. The labelled peptides were cleaved and purified as described above. The peptides were purified on a Waters preparative HPLC using a reverse-phase Waters C18 semi-preparative column with a gradient of 5:60 % ACN in TDW. The column was equilibrated with 5% Acetonitrile (ACN) for 5 minutes, followed by a linear increase from 5% to 60% ACN over 40 minutes. The identity and purity of the peptides were determined using MALDI-TOF mass spectrometry and analytical HPLC.

### Flow Cytometry Analysis of Peptide-Bacteria Binding

The bacteria were recovered overnight by inoculating a colony from a plate into fresh Luria broth (LB) medium in a shaking incubator at 37°C. After recovery, 100 mL of the bacteria were transferred into fresh 5ml LB to grow until they reached OD600_nm_<0.5. The cultures were centrifuged (4000RPM, 4°C), and their pellet was washed with PBS 3 times and diluted to OD600_nm_=0.1. The Fluorescein-labelled peptides were dissolved in fresh PBS and filtered using 0.45 µ and 0.2 µ filters and diluted to 0.1μM to 25 μM concentrations. The bacteria and peptides were incubated for 20 minutes and washed 2 times with PBS by 2500RMP centrifugation for 30 seconds. Each sample was transferred to a 96-well plate for Flow Cytometry analysis.

For the Flow cytometry, we used the Aurora Cytek. Data for the following parameters were collected and used for setting gates in the cell sorter: FSC – Forward Scatter, SSC – Side Scatter, 488nm intensity – close to the excitation wavelength of FAM. The data recorded was for 5000 events. The first gate was to differentiate between bacterial cells and debris in the sample using the FCS and SSC as the gating axis. Bacterial cells have a natural autofluorescence of up to 1.1×10^3^ A.U. (Fig. S1), so the next gate was from 1.1×10^3^ A.U. up to the intensity limit of the machine 2×10^6^ A.U., on the 488nm absorbance axis (x-axis). All measurements were performed with at least 3 independent replicates. Afterwards, the data were analyzed using an R software script.

### Biolayer Interferometry (BLI) Binding Assays

BLI experiments were performed using Gator Prime (GatorBio). 5 ug of recombinant His-tagged InlA (Biorbyt, orb1674721) was immobilized onto Ni-NTA biosensors. An initial baseline was established in PBSX1 (pH 7.4), ionic strength 154.85 mM. For the association phase, the functionalized biosensors were transferred into solutions containing the E-Cad1 peptides at varying concentrations (10 to 100 µM for E-Cad1 11-31 and E-Cad1 22-31, and 10 to 20 µM for E-Cad1 15-24) for 300 seconds. The sensors were subsequently moved into peptide-free PBS for a dissociation phase of 600 seconds. All measurements were performed at 25 °C with continuous shaking.

Sensorgrams were reference-subtracted to correct for baseline drift and bulk refractive index shifts. Data analysis was performed using the GatorOne software (GatorBio). The curves were fit to a local 1:1 kinetic binding model to determine the apparent equilibrium dissociation constants (K_D_). The association and dissociation phases were analyzed for 100 seconds of the association and 600 seconds of dissociation for E-Cad1 11-31, and 100 seconds of association and 300 seconds of dissociation for E-Cad1 15-24. The reported K_D_ values are calculated as the mean ± standard deviation across the evaluated concentration ranges.

### Gold working Electrode Preparation and modification

Bare gold electrodes (**AuE**) were manually polished using alumina powder (0.05 µm) suspensions in TDW (Fig. 5. Step I), followed by extensive washing with deionized water to remove residual abrasive particles. Following this, the electrodes were washed sequentially in EtOH and TDW to ensure thorough removal of any remaining impurities before peptide functionalization. Following the functionalization step, the electrode was modified with a 50 μM peptide solution in TDW, and the electrode was incubated for 10min at 37 °C (Fig 4. Step II). After the modification, the electrodes were rinsed with TDW and EtOH. After the modification, the electrode was exposed to the solution containing the bacteria in different concentrations suspended in PBS and incubated for 10 min. at 37°C (Fig4. Step III).

### Electrochemical Measurements

The Electrochemical impedance spectroscopy (EIS) measurements were performed in a standard three-electrode electrochemical cell setup. The AC potential source is provided by a BioLogic SAS SP-300 potentiostat under single sine AC excitation at a potential of 0.21 V with 10 mV amplitude in the frequency range from 100 kHz to 0.1 Hz. The electrochemical cell comprises a reference electrode, Ag/AgCl, **AuE** working electrode (2 mm), and a Pt billet as a counter electrode. The measurements were performed in [Fe(CN)6]^3−^/[Fe(CN)6]^4−^] 5mM and KCl 100 mM solution. The results were fitted to the equivalent circuit of RS[(R_CT_W)∥Q]. RS is the resistance of the solution, R_CT_ is the charge-transfer resistance of the layer, Q is the constant phase element, and W is the 120 Warburg diffusion element. The normalized R_CT_ is calculated as the ratio between the R_CT_ after and before the **AuE** exposure to bacteria. Molecular density acquired by using reductive desorption analysis, which was performed for the determination of surface coverage of the LPA coupled E-Cad1 15-24 on **AuE**. This characterization was done by CV in KOH [0.5 M] electrolyte with degassing by N2 for 5 min before the analysis. The reductive desorption electrochemical sweep was between −0.5 and −1.4 V with a scan rate of 100mV/s.

### Surface Characterization

Surface characterizations, XPS, CPD, and VASE, were performed on a Au (111) layer (100 nm) on top of a Cr layer (10 nm), which was vapor deposited on the substrate of an n-type Si wafer (100). The bare gold surfaces were washed with absolute EtOH before and after 20 min of cleaning using Ultraviolet Ozone Cleaning Systems (UVOCS Inc). VASE measurements were performed with an α-SE ellipsometer (J.A. Woollam Co.) at a silicon Brewster angle of 75°, and the results were fitted with the Cauchy model for the organic layer. PS measurements were performed using an Axis Supra+ spectrometer (Kratos Analytical Ltd., Manchester, U.K.) with an Al Kα monochromatic X-ray source (1486.7eV). The XPS spectra were acquired with a takeoff angle of 90° (normal to analyzer); the vacuum condition in the chamber was 1.9 nTorr. High-resolution XPS spectra were acquired with a pass energy of 20 eV and a step size of 0.1 eV. The binding energies were calibrated according to the C 1s peak position (285.0 eV). Data were collected and analyzed by using Casa XPS (Casa Software Ltd.) and the Vision data processing program (Kratos Analytical Ltd.).

### Microscopy measurements

Peptide-functionalized Au(111) layers (100 nm), deposited on a Cr adhesion layer (10 nm) over n-type Si (100) substrates, were incubated separately with three bacterial strains at a concentration of 10 CFU /mL under the same conditions used for the electrochemical sensing protocol. Following incubation, the electrodes were rinsed with PBS to remove unbound cells and stained with Nile Red fluorescence dye (1 µg/mL in PBS, 10 min, 37°C) as to visualize bacterial membranes. Excess dye was removed by additional PBS washes. The fluorescence microscopy was performed using a Zeiss Axioskop2 microscope equipped with an XBO75 light source. Images were captured using DinoCapture 2.0 software. Fluorescence was observed using filter sets appropriate for Nile Red (excitation ∼552 nm, emission ∼636 nm). All imaging was performed at 100× magnification, with a scale bar of 100 µm included in each image.

### Biosensor exposure to Bacteria

The bacterial cells were grown overnight in LB at 37°C. The bacterial cell suspension was diluted 1/50 in fresh LB and grown for an additional 1-2 h. The bacterial pellets were washed twice with PBS and separated by centrifugation (8000 rpm for 8 min, 4°C) and diluted to an optical density (OD) of 0.1 at 600_nm_, corresponding with 10^8^ CFU/ml. The desired concentration was obtained by serial dilution in PBS and applied in a volume of 1 ml onto the AuE-E-Cad1. Two additional electrodes were maintained in PBS buffer as controls, showing minimal change in their R_CT_ values. The limit of detection (LOD) was calculated using the standard 3σ method, where 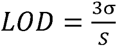 with σ representing the standard deviation of replicate measurements (n=5) at the lowest bacterial concentration (1 CFU/mL, σ = 0.05) and S representing the slope of the calibration curve. This approach is widely accepted for biosensor validation, as it defines the minimum analyte concentration that produces a signal statistically distinguishable from baseline noise with 99% confidence.

## Acknowledgements

This research is supported by the National Research Foundation, Prime Minister’s Office, Singapore, under its Campus for Research Excellence and Technological Enterprise (CREATE) programme, through the Cellular Agriculture Programme (CellAg) No:370184512. AF thanks The Minerva Center for Bio-Hybrid complex systems and the Saerree K. and Louis P. Fiedler Chair in Chemistry. S. Y. thanks the Benjamin H. Birstein Chair in Chemistry.

