## Supporting information for "Rapid Electrochemical Biosensing of *Listeria monocytogenes* Using Rationally Designed Host-Pathogen Interface Peptides"

**Table S1: Sequences and masses of designed peptides**

| **Peptide name** | **Sequence** | **Calculated Mw of peptides (g/mol)** | **Mw +** **FAM** | **Mw +** **LPA** |
| --- | --- | --- | --- | --- |
| E-Cad1 11-31 | _11_ENEKGPFPKNLVQIKSNKDKE_31_ | 2441.748 | 2801.06 | 2631.06 |
| E-Cad1 15-24 | _15_GPFPKNLVQI_24_ | 1111.334 | 1469.65 | 1300.64 |
| E-Cad1 22-31 | _22_VQIKSNKDKE_31_ | 1187.343 | 1545.66 | 1375.66 |

**FACS binding assay controls – bacteria only**

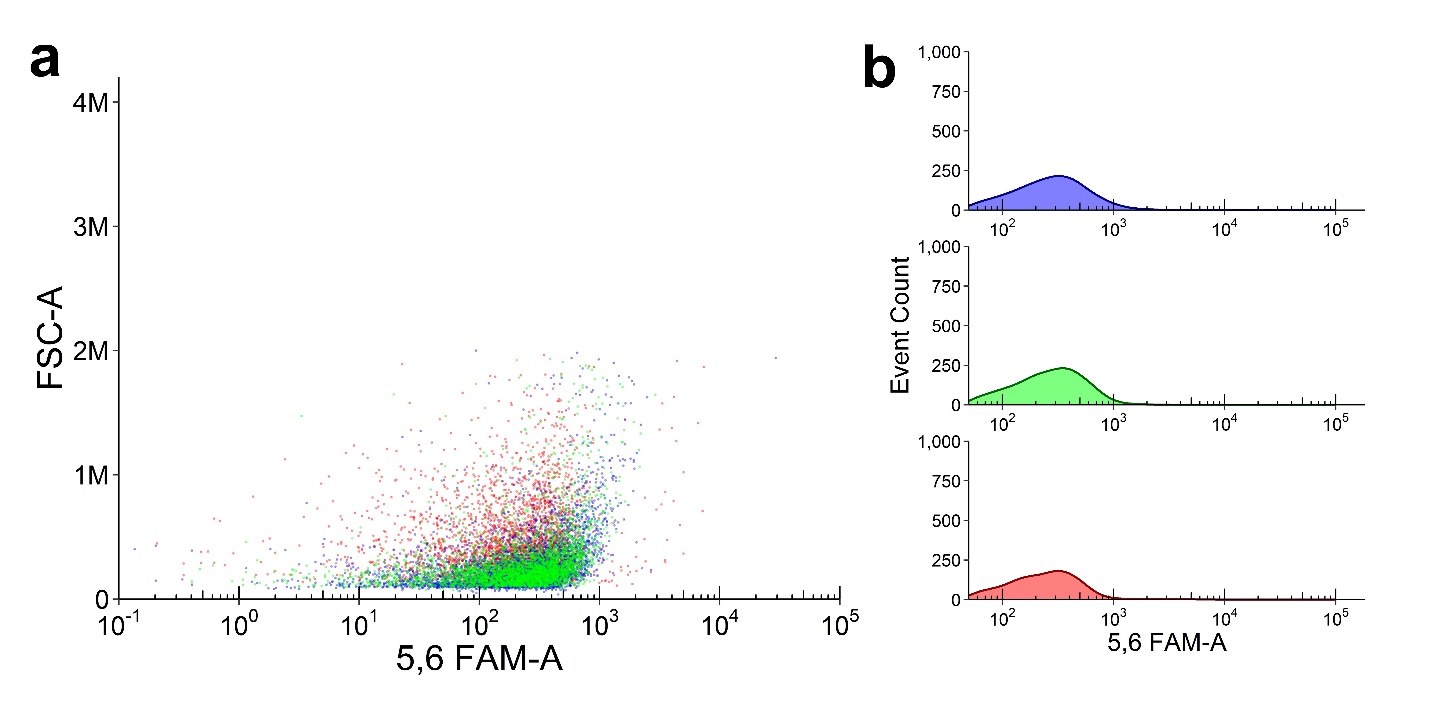

**Figure S1.** Autofluorescence of *LM* (Green) compared to *E. coli* (Blue) and *MRSA* (Red) without peptides analyzed using flow cytometry. **a.** Dot plots of fluorescence intensity against FCS showing all bacteria are in the lower fluorescence range. **b** Event count histograms of fluorescence intensity for *LM*, *E. coli and MRSA*.

**Reductive desorption-derived peptide surface number density on gold**

We were able to calculate the number of Peptides and surface concentration of molecules on AuE. Based on the Equation, Peptide =(Q*C)/(A*n), the number of peptides on the AuE surface, Q is the charge transfer (calculated by the integral of the CV peak), an average of 3.11×10^-7^ coulomb, C is the coulomb constant, and A is the area of the electrode 3.14 mm^2^. Since each peptide molecule has two S-Au bonds, the reduction of each molecule is associated with 2 electrons, n = 2 (Figure S1).

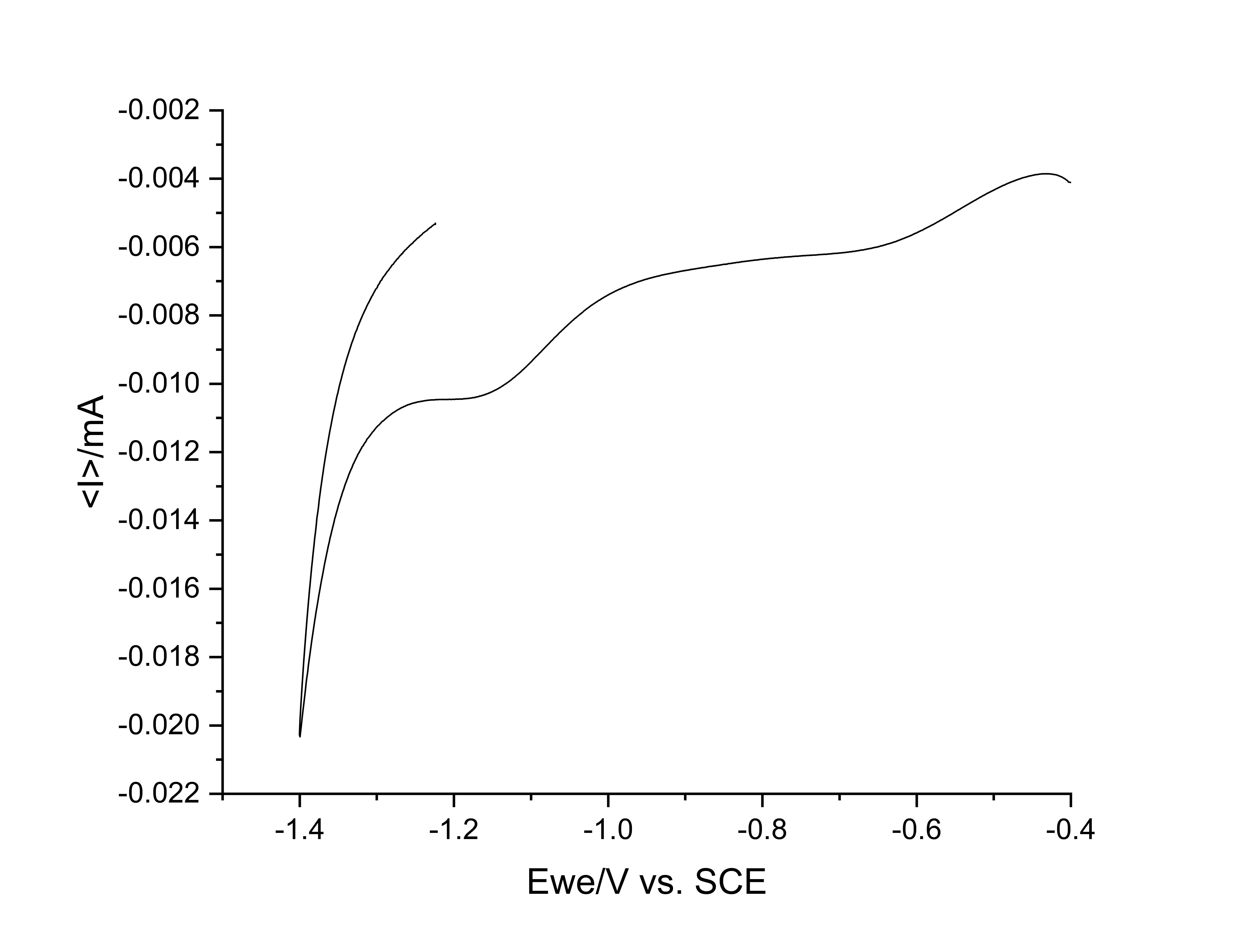

**Figure S2.** Reductive desorption analysis based on CV data (See Methods).

**Surface analysis by XPS**

The thickness of organic thin films deposited on substrates was quantified by X-ray photoelectron spectroscopy (XPS) using a substrate signal attenuation model. This approach utilizes the exponential decrease in substrate core-level photoelectron intensity as a function of overlayer thickness due to inelastic scattering, following the Beer–Lambert law. For a film of thickness, the intensity of the substrate signal is $I_{\text{Overlayer}}$, and the bare substrate intensity is$I_{\text{sub}}$, $\theta$ is the mission take-off angle to the surface normal. All spectra were background corrected and peak areas integrated with a calibrated transmission function and appropriate sensitivity factors. Values of $\lambda$ were taken from TPP-2M or literature tables and matched to the kinetic energy of the observed core levels in the respective matrix (Table S2, Figure S2).

**Table S2: XPS C1s and Au 4f signals and peptide thickness calculation**

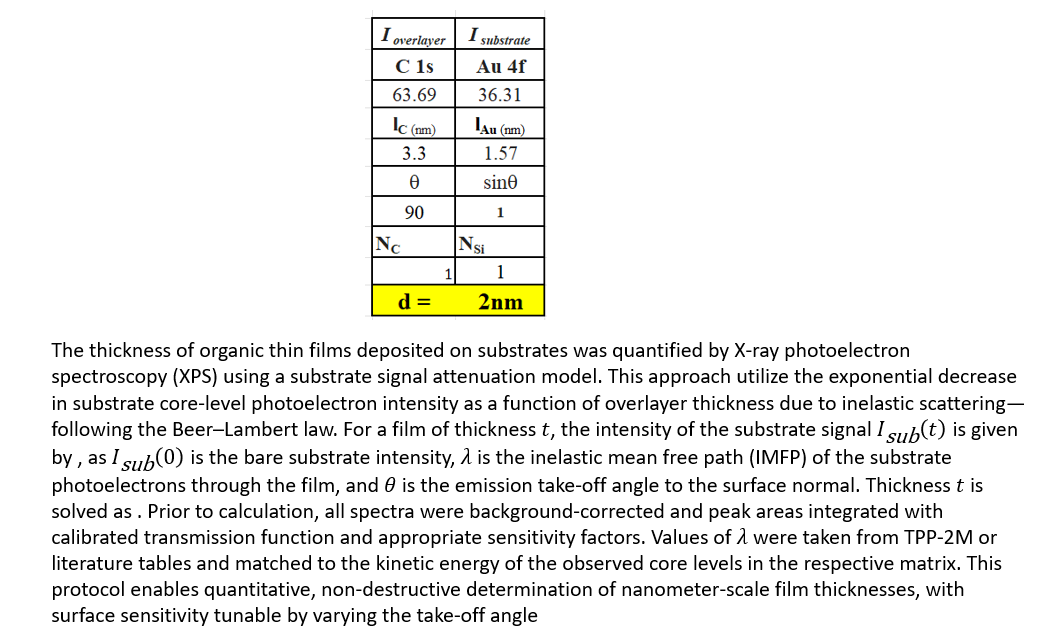

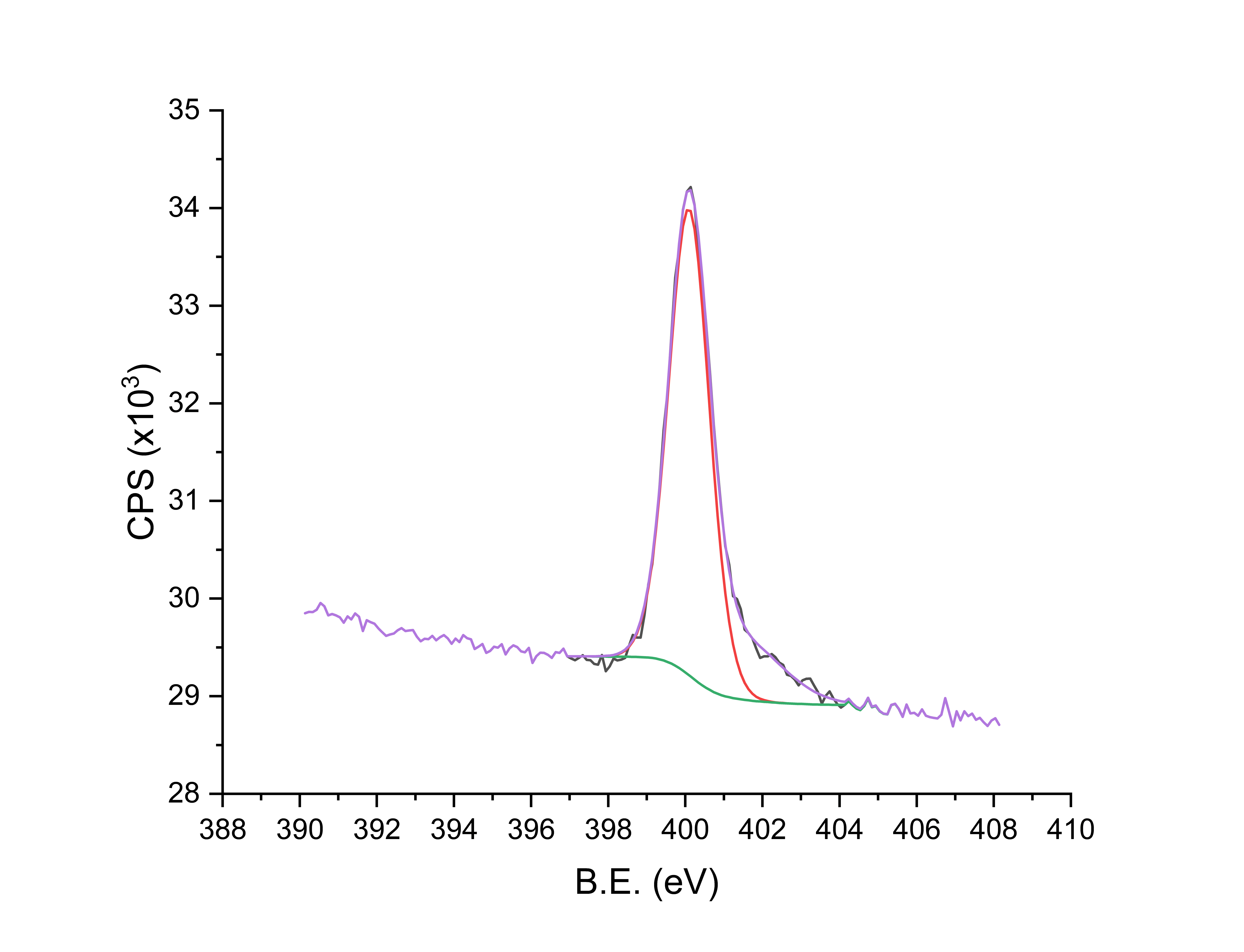

**Figure S2**. XPS N1s signal; raw data is represented as CPS (black), background (green), N-C bond (orange), and deconvoluted envelope (purple).

The distinctive peak at 401 eV characteristic of the amide bond supports the modification of the AuE with the peptide.

**Fluorescence microscopy confirms LM binding to the peptide-grafted surfaces**

To visualize the bacterial detection of *LM* by the E-Cad1 15-24 peptide, gold plates were functionalized with the peptide and incubated with bacterial cultures of *LM*, *MRSA,* and *E. coli* under the same conditions used for electrochemical sensing. Following incubation, electrodes were stained with Nile Red to label bacterial membranes and imaged using fluorescence microscopy. The fluorescence images revealed clear and localized binding of *LM* to the peptide-coated gold surfaces with no significant fluorescence for images of the control bacterial strains. Nile Red staining produced distinct fluorescent signals corresponding to bacterial cells, confirming successful interaction with the immobilized E-Cad1 15-24 peptide. (Fig. S3)

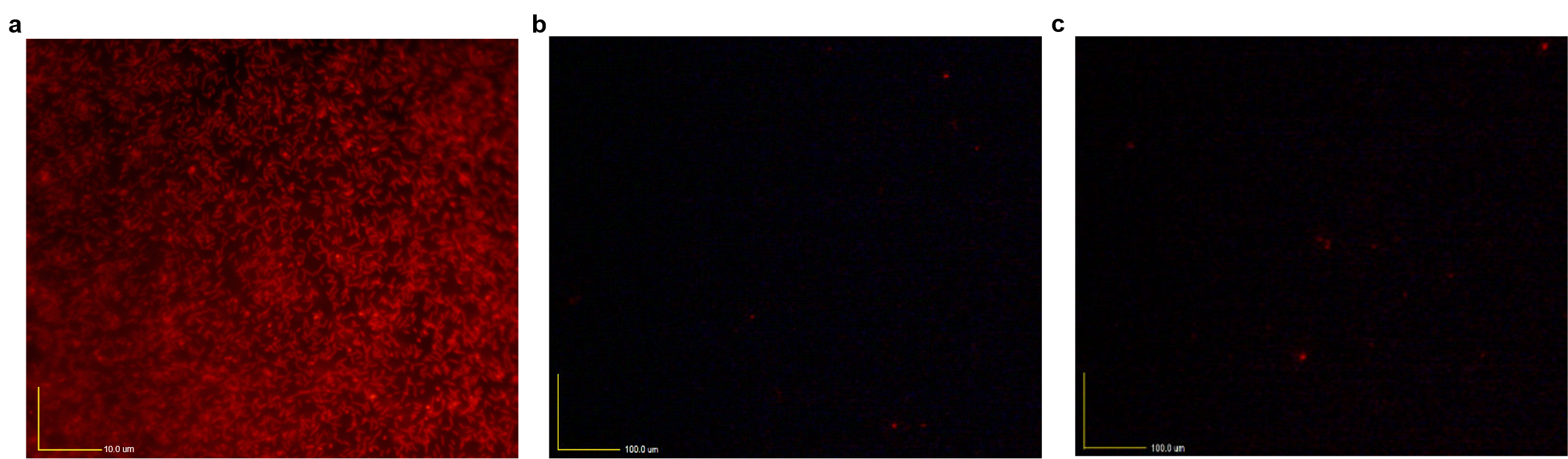

**Figure S3.** Fluorescence microscopy images of E-Cad1 15-24-LPA functionalized **AuE** and incubated with different bacterial strains, followed by Nile Red staining. Images were acquired using a fluorescence microscope at 100× magnification. Scale bar: 100 µm. **a** *LM* shows strong and localized fluorescence, indicating specific binding to the peptide-coated surface. **b** *E. coli* shows sparse and faint fluorescence, suggesting minimal non-specific interaction. **c** *MRSA* shows few and weak fluorescent signals, indicating limited or non-specific binding.
